# FLInt 2.0: Robust and customizable single shot integration in *C. elegans*

**DOI:** 10.64898/2026.01.12.699147

**Authors:** Nawaphat Malaiwong, Porhathai Malaiwong, Chloe Kim, Michael O’Donnell

## Abstract

Transgenesis in *Caenorhabditis elegans* has revolutionized biological research by enabling the precise control of expression of both endogenous and exogenous genes. The FLInt (<u>F</u>luorescent <u>L</u>andmark <u>Int</u>erference) method was developed to accelerate site-specific integration of transgenes, but persistent false-positive events during screening have emerged as a rate-limiting step. Here, we present an alternative FLInt strategy, FLInt 2.0, that reduces false positives by biasing Cas9 cutting of the tdTomato and Cbr *unc-119*(+) safe-harbor locus, adjacent to the untranslated region (3’ UTR). The design preserves fluorescence during non-integrative repair events, such that only true integration of an extrachromosomal array abolishes tdTomato expression. We demonstrate that this targeted approach maintains high integration efficiency while significantly decreasing the proportion false positives observed in F1 progeny. Molecular and transmission analyses confirm that non-fluorescent F2 animals reliably represent stably integrated multi-copy transgenic lines, which can be tailored to desired expression levels using a simple subsequent Cas9 targeting approach. By exploiting coding-frame geometry to discriminate integration events from repair-induced artifacts, this method streamlines the identification of the true integrants, reducing labor-intensive screening and increasing experimental throughput. Our strategy provides a robust, visually guided, and efficient refinement of FLInt, offering a generalizable framework for improving site-specific transgene integration in *C. elegans*.

## Introduction

Integration of exogenous transgenes advanced the utility of *Caenorhabditis elegans* as a model organism by enabling genetic manipulation, protein localization studies, and targeted modulation of cellular processes (Nance and Frøkjær-Jensen 2019). Transgenes are typically introduced via gonadal microinjection of plasmid DNA, which assembles into large extrachromosomal arrays through *de novo* recombination (Stinchcomb et al. 1985; Mello et al. 1991). While these arrays allow robust expression—including of heterologous genes—they are not chromosomally integrated, resulting in unstable inheritance and mosaicism. To address this, a variety of integration techniques have been developed to ensure stable transmission and more uniform expression. Integration strategies vary in terms of copy number, site specificity, and procedural complexity. Single-copy insertion methods via transposon elements e.g. MosSCI and MiniMos (Frøkjaer-Jensen et al. 2008; Frøkjær-Jensen et al. 2014), FLP recombinase (Nonet 2020; Nonet 2023), PhiC31 integrase (Yang et al. 2022), and CRISPR/Cas9 (Norris et al. 2015; Philip et al. 2019; Silva-García et al. 2019; Stevenson et al. 2020) offer precise targeting and reproducible outcomes. However, in many cases multi-copy transgene insertion is preferable due to increased expression levels or to combinatorially express multiple transgenes.

Multi-copy integration can be achieved in both targeted and random manners using CRISPR (Yoshina et al. 2016; El Mouridi et al. 2022; Yoshina and Mitani 2022; Malaiwong et al. 2023; Yanagi and Lehrbach 2024), UV or gamma irradiation (Mello and Fire 1995; Evans 2006; Mariol et al. 2013), PhiC31 integrase (Rich et al. 2025), and miniSOG (Noma and Jin 2018), respectively. Random insertion approaches via UV and miniSOG carry risks of off-target effects and often require labor intensive outcrossing and/or mapping. Among targeted approaches, a major portion of effort is often spent isolating rare integration events from a large population of false positives, as extrachromosomal array formation occurs at a higher frequency than genomic integration (Malaiwong et al. 2023). Recently established strategies to enrich for true integrants include *unc-119* phenotypic rescue (El Mouridi et al. 2022; Yanagi and Lehrbach 2024), drug resistance (Dickinson et al. 2015; Stevenson et al. 2023; Rich et al. 2025), and fluorescent protein expression (Malaiwong et al. 2023). These methods require either cloning of transgenes into specialized targeting vectors or may yield false positives requiring screening, either of which may serve as a barrier for uptake of these approaches for some of the *C. elegans* research community. FLInt (Fluorescent Landmark Interference) was originally developed as a simplified multi-copy integration approach that leverages CRISPR/Cas9-mediated excision of a defined locus bearing a fluorescent marker – a “safe harbor” site – and required no specialized reagents or cloning (Malaiwong et al. 2023). A large number of strains bearing these safe harbor sites exist (Frøkjær-Jensen et al. 2014) and by targeting the encoded tdTomato::H2B using a crRNA, FLInt enables detection of CRISPR activation in F1 and integrated transgenes in the F2 generation through loss of red fluorescence. However, despite its streamlined one-shot injection protocol, FLInt still demands substantial effort to identify true integrants due to numerous false positives arising from non-integrative mutagenic repair events. This rate-limiting step underscores the ongoing trade-off between procedural simplicity and the screening workload required for accurate identification.

Here we present an improved single shot approach for the generation of multi-copy integrated transgenes which we have termed “FLInt 2.0”. FLInt 2.0 enables a simple visual method to distinguish true integrants from mutagenic non-integrants. We show that by introducing double strand breaks near the junction of the fluorescent protein coding domain and a 3’ UTR, most non-integrative mutagenic events due to repair do not interfere with fluorescence. In contrast, integration of multi-copy arrays disrupts this junction and results in complete elimination of fluorescent protein expression. As a result, the fluorescent animals can be easily excluded during screening. We show that this optimized approach results in nearly 100% accurate identification of true integrants and requires a minimal screening effort that is comparable to extrachromosomal array generation. We further show that the expression level of integrated multi-copy arrays using this method can be customized via a simple subsequent round of CRISPR-mediated excision using sequences common to most cloning vectors. This flexible, low-cost and streamlined method should enable the generation of integrated transgenes with few barriers compared to extrachromosomal arrays.

## Materials and methods

### Animal husbandry

*C. elegans* strains were maintained using standard procedures (Stiernagle 2006).Wild-type (N2) and parental integration strains were obtained from the Caenorhabditis Genetics Center: EG7837 (*oxTi712* [*eft-3*p::tdTomato::H2B::*unc-54* 3’UTR + Cbr-*unc-119*(+) I]; *unc-119*(*ed3*) III), EG7944 (*oxTi553* [*eft-3*p::tdTomato::H2B::*unc-54* 3’UTR + Cbr-*unc-119*(+) V]); *unc-119*(*ed3*) III), and were cultured at 20 °C prior to microinjection. Following injection, animals were maintained at 25 °C to accelerate developmental rates. As noted by CGC, tdTomato-expressing strains often appear sick when incubated at 25 °C, likely due to the ubiquitous expression of the fluorescent marker.

### crRNA designs

crRNAs for CRISPR/Cas9 were designed using the “Design CRISPR Guides” tool on Benchling and evaluated for efficiency and specificity with the IDT CRISPR-Cas9 guide RNA checker. The tdTomato-targeting crRNA (5’-AGATGGTCTTGAACTCCACC-3’) was modified from the original report (Malaiwong et al. 2023), with additional guides designed against the 3′ region of tdTomato (5’-CGTCCATGCCGTACAGGAAC-3’) and the tdTomato linker (5′-CTCCTCCGAGGACAACAACA-3′). A crRNA targeting the H2B (5’-ATGCTAACTAGTTTACTTGC-3’) was designed to cut at the H2B::3′UTR junction with no predicted genomic off-targets, and another crRNA was designed against the *C. briggsae unc-119* (5′-TTATGCATCATATGAGTAGT-3′). All crRNAs were verified to lack significant complementarity to the *C. elegans* genome or other fluorescent reporters, minimizing the risk of off-target effects.

### Molecular techniques

The DNA fragments were amplified from the plasmid templates using Phusion™ High-Fidelity PCR Master Mix with HF Buffer (NEB) or KOD Hot Start DNA Polymerase (Sigma-Aldrich, cat. no. 71086-3). The plasmids were constructed using the NEBuilder® HiFi DNA Assembly Master Mix (NEB). The solution was then transformed into the TSS competent cells. The plasmids were extracted from bacteria cultures using Zyppy Plasmid Miniprep Kit (ZymoResearch, #D4020). The PCR for genotyping was done using OneTaq® 2X Master Mix with Standard Buffer (NEB, #M0482L).

### Microinjection

Worm injections were performed following standard protocols (Evans TC. 2006). To improve the animal’s viability and brood size, a paint brush was used for transferring worms during microinjection (Gibney et al. 2023). 10-15 injected P0 worms were transferred together onto 6 cm NGM/OP50 plates and cultured at 25°C.

### Hygromycin treatment

Hygromycin stock solution (5 mg/mL) was prepared by diluting Hygromycin B (50 mg/mL) (ThermoFisher Scientific, #10687010) with milliQ water and stored at 4°C. The stock solution was applied by top spreading (500 uL per 10-mL agar plate) and bench dried for 20-30 minutes. As hygromycin is harmful, the PPE should be applied during this step.

### Visual screening of candidates

Candidates for array integration were typically identified ∼6 days post-injection. Hygromycin-treated plates containing F1–F2 progeny were screened under a fluorescence stereomicroscope, and candidate integrants were selected based on complete loss of tdTomato::H2B fluorescence. Selected animals were transferred to small NGM/OP50 plates for F3 screening to identify homozygous integrants. At this stage, dim residual tdTomato signal in F2 animals—likely arising from specific INDEL alleles—can increase false-positive selection; therefore, worms exhibiting partial or dim red fluorescence were avoided. In our experiments, candidates were initially selected solely based on loss of tdTomato::H2B without considering array expression (indicated by pharyngeal *myo-2*p::GFP, Vázquez-Manrique et al. 2010), and we observed that tdTomato-negative animals frequently retained array expression. Final verification of integrated lines was performed in the F3 generation, using array marker expression as a visual confirmation.

### Quantification of transgene copy number

Genomic DNA was prepared from *C. elegans* strains carrying integrated GFP constructs. For each sample, 10 adult worms were lysed in 20 µL worm lysis buffer (50 mM KCl, 10 mM Tris–HCl pH 8.3, 2.5 mM MgCl₂, 0.45% NP-40, 0.45% Tween-20, 0.01% gelatin, and 60 µg/mL proteinase K). Lysates were incubated at 65 °C for 1 h and heat-inactivated at 95 °C for 15 min, then stored at −20 °C. Quantitative PCR (qPCR) was performed using 2 µL of worm lysate, corresponding to the DNA content of a single worm, as template in a 10 µL reaction with iTaq Universal SYBR Green Supermix (Bio-Rad) on a Bio-Rad CFX Opus 96 Real-time PCR System. Primer pairs were designed to amplify GFP (Forward primer: 5’-ATGAGTAAAGGAGAAGAACTTTTCACTGGAGTTGTC-3’ and Reverse primer: 5’-CCTTCAAACTTGACTTCAGCACCTG-3’) and the single-copy endogenous *act-1* gene (Forward primer: 5’-TTGCCGCTCTTGTTGTAGACAA-3’ and Reverse primer: 5-ATTGGGTACTTGAGGGTAAGGA-3’) which served as the reference locus. Thermal cycling was performed with an initial denaturation at 95 °C for 10 min, followed by 40 cycles of 95 °C for 15 s and 60 °C for 1 min. Ct values for each strain were taken as the median of 3 technical triplicates. Transgene copy number relative to *act-1* was estimated by the comparative Ct method assuming 100% PCR efficiency. For each strain, ΔCt_strain_ was calculated as Ct_GFP_−Ct_act−1_. ΔΔCt_strain_ was calculated as ΔCt_strain_ - ΔCt_single-copy,_ where ΔCt_single-copy_ represented the single-copy reference strain, respectively. Relative copy number was determined as 2^-ΔΔCt(strain)^. Strains analyzed included three integration strategies: MOY258–MOY262 (original FLInt method), MOY263–MOY267 (FLInt with tdTomato linker cutting site), and MOY226–MOY235 (FLInt with 3′ tdTomato cutting site).

### Statistical analysis

Statistical analyses were performed using an unpaired two-tailed *t*-test, as specified in the figure legends.

### Microscopy

Fluorescence microscopy was performed using a Leica DMi8 inverted fluorescence microscope equipped with a K5 Scientific sCMOS camera. Images were acquired using LAS X software. Adult worms were mounted on 2% agarose pads containing 10 mM tetramisole for immobilization. Single-plane captures and Z-stack images were acquired and processed using the Leica THUNDER Imager system.

### Quantification of flourescence

#### tdTomato

To measure tdTomato expression levels, fluorescence intensity in F1 progeny was quantified at the L4/young adult stage following injection with the FLInt mixture. Three days post-injection, F1 animals were screened for the *myo-2*p::GFP array marker. Array-positive worms were mounted on 2% agar pads, immobilized with 10 mM tetramisole, and imaged. Red fluorescence was captured using a 4x air objective (17% FIM power, 100 ms exposure). tdTomato intensity was measured in ImageJ by defining a polygonal ROI tracing the worm’s outline. Background fluorescence was measured from an adjacent region outside the worm and subtracted from the ROI signal. Final fluorescence values (A.U.) were calculated as *Intensity(ROI) – Intensity(background)*. tdTomato intensities from P0, F1, and F2 animals generated using FLInt1.0 and FLInt2.0 plotted and compared. To compare F1 fluorescence phenotypes, raw tdTomato fluorescence intensity values measured in F1 progeny were classified into three categories based on predefined intensity thresholds: OFF (≤120), intermediate (120–180), and ON (≥180). For each experimental condition, the proportion of animals in each category was calculated and expressed as a percentage of the total population. These percentages were visualized using stacked bar plots. These distributions were compared across different FLInt injection mixtures.

#### Pharyngeal GFP

To compare the pharyngeal markers in tuning transgene copy number strategy, the intensity of the pharyngeal marker *myo-2*p::GFP (A.U.) was quantified by imaging young adult transgenic worms mounted on glass slides using a 4X objective. The GFP fluorescent was excited by 17% laser intensity, 10ms exposure time. Green fluorescence was measured from the metacorpus region of the pharynx using ImageJ. A circular ROI was drawn over the metacorpus to obtain the signal intensity.

#### CAN cell

The intensity of CAN::mCherry was quantified by imaging young adult transgenic worms mounted on glass slides using a 20X objective lens. Red fluorescence was measured from the CAN cell region using ImageJ. A circular region of interest (ROI) was drawn over the CAN cell to obtain the signal intensity.

### Next generation sequencing

NGS was performed to quantify INDEL variants generated at different developmental stages following CRISPR activation. CRISPR-edited animals were produced by microinjecting a CRISPR mix targeting the H2B region of the tdTomato landing site together with a co-injection marker (3 ng/µL *myo-2*p::GFP) and 97 ng/µL DNA ladder (Invitrogen). Injected worms were maintained at 25 °C. At 36 h post-injection, array-positive L2–L3 larvae were identified by marker expression and lysed either individually or as pooled samples. At the adult stage, array-positive F1 progeny were selected and processed as pooled lysates (15 worms per tube) or as single-worm lysates. A subset of array-positive F1 animals was propagated to generate F2 progeny. Adult F2 animals retaining the array were collected either as pooled samples (three independent lines, five worms per line) or as individual worms. All lysates were stored at −20 °C prior to PCR. NGS amplicons were generated using KOD Hot Start DNA Polymerase (Sigma-Aldrich, cat. no. 71086-3) in 50 µL reactions containing 2 µL of worm lysate with locus-specific primers (Table 1). Reactions were purified using the QIAquick PCR Purification Kit (QIAGEN, cat. no. 28106) and processed by Quintara Biosciences via 250bp paired-end Illumina sequencing.

**Table 1.**
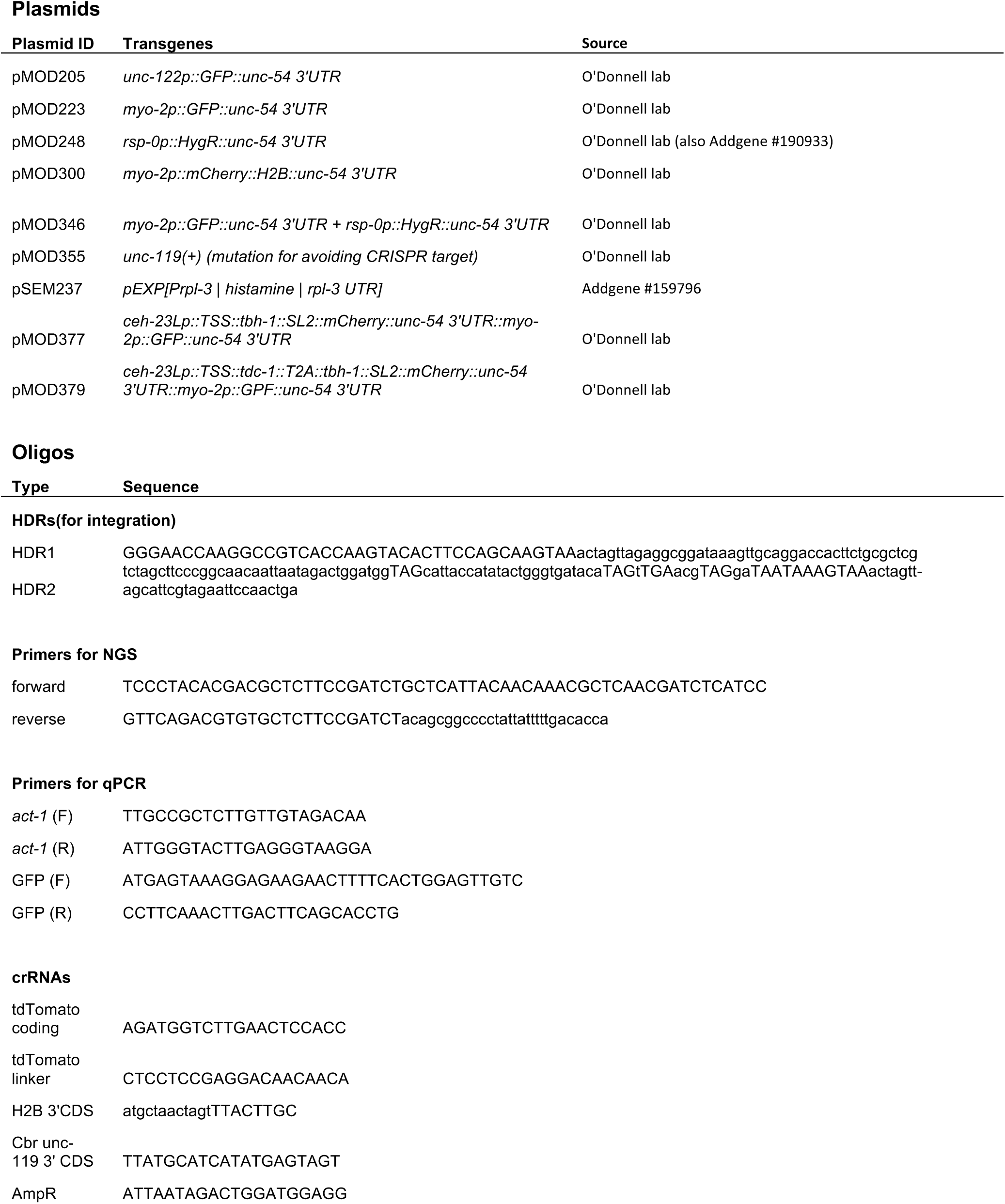
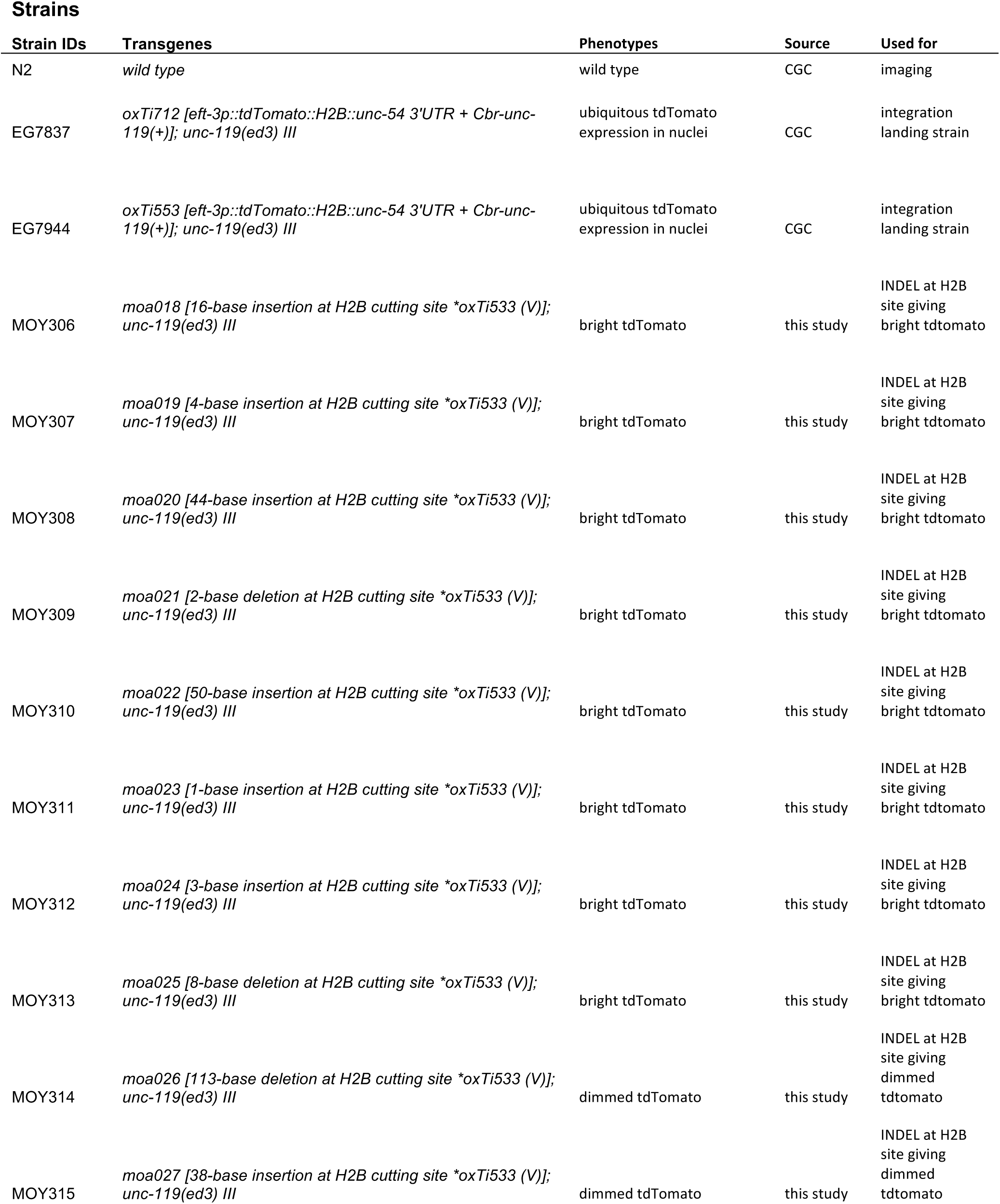
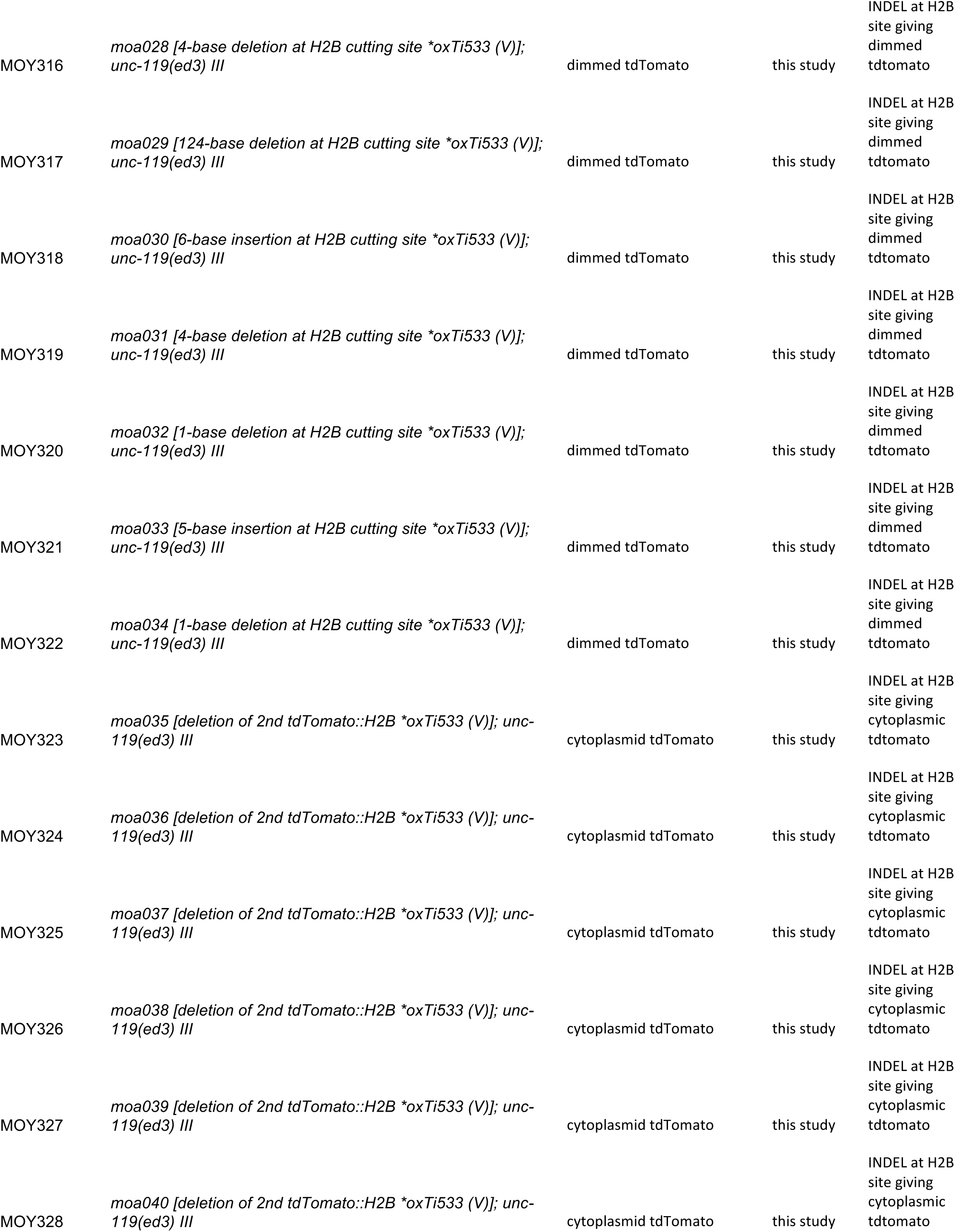
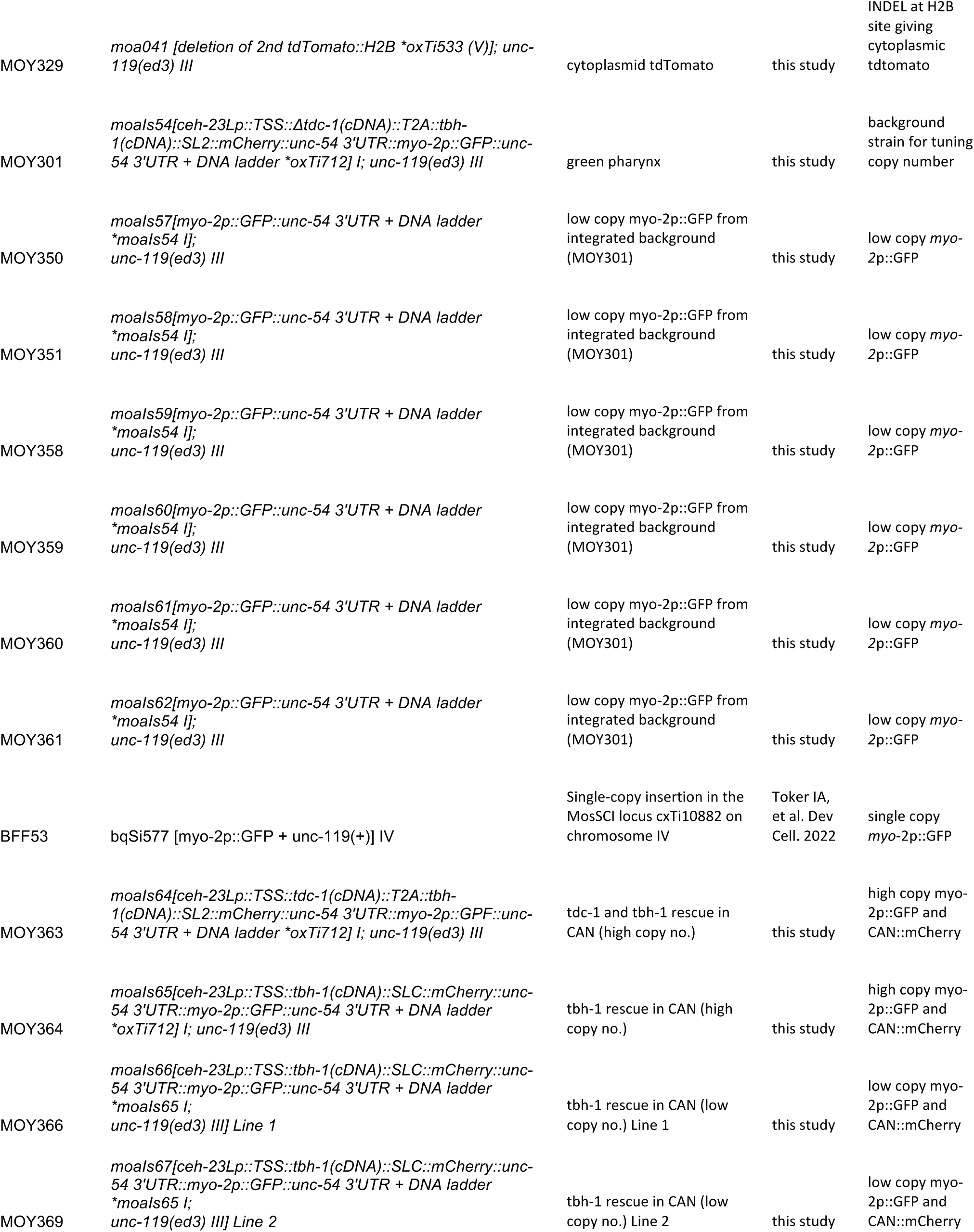
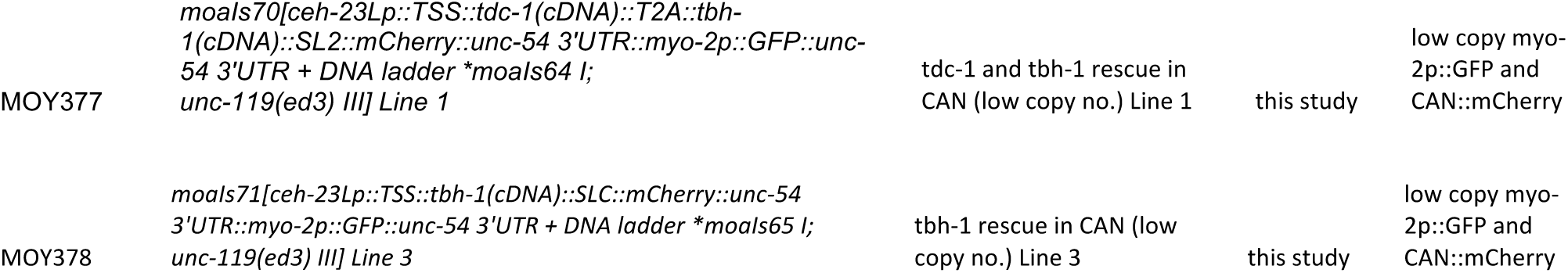
Reagents.

### Analysis of NGS amplicon sequences

INDEL variants were extracted, quantified, and compared across non-edited controls, L3 F1 larvae, adult F1 animals, and adult F2 animals. Raw reads were filtered above Phred quality >=30 using Fastp. Sequence depth was between 30K-80K reads per sample after filtering. Filtered reads were then run through a custom script to count unique sequences. Initial observations using unedited worm sequence reads indicated PCR or sequencing errors occurred at a rate of < 1%, so this was used as a cutoff for quantification of edited alleles. All alleles above this threshold were then aligned using a custom python script, and alignments were manually inspected to confirm that the majority of sequence variants remaining occurred at or near the intended cut site.

## Results

### Specific CRISPR-mediated cutting of safe harbor sites preserve fluorescence in the absence of integration

Sequencing of previously generated integrated transgenic worms using the FLInt 1.0 method indicated that there were principally 2 classes of animals lacking fluorescence after injection: those bearing small insertions or deletions (INDELs) and those containing large, multi-copy array insertions (Malaiwong et al. 2023). The former class represented the majority of isolated worms, and are thus considered “false positives”, representing increased experimental effort and screening. Because these classes were initially indistinguishable, we reasoned that maintaining fluorescence in animals bearing small indels would improve selection accuracy by ensuring that only true integrants—which lose the tdTomato reporter through extrachromosomal array insertion—would become non-fluorescent only when homozygous in the F2 generation. To test this idea, we identified CRISPR target sites that are predicted to generate different fluorophore expression levels depending on the mechanism of gene editing, focusing on a collection of existing chromosomally integrated single copy transgene strains that express ubiquitous nuclear-localized tdTomato (*eft-3*p::tdTomato::H2B::*unc-54* UTR; Cbr-*unc-119*(+)), enabling efficient visual screening. We targeted sites that are predicted to preserve an intact promoter–coding sequence–UTR configuration after the generation of small indels (Fig 1A, Frøkjær-Jensen et al. 2014).

**Figure 1.**
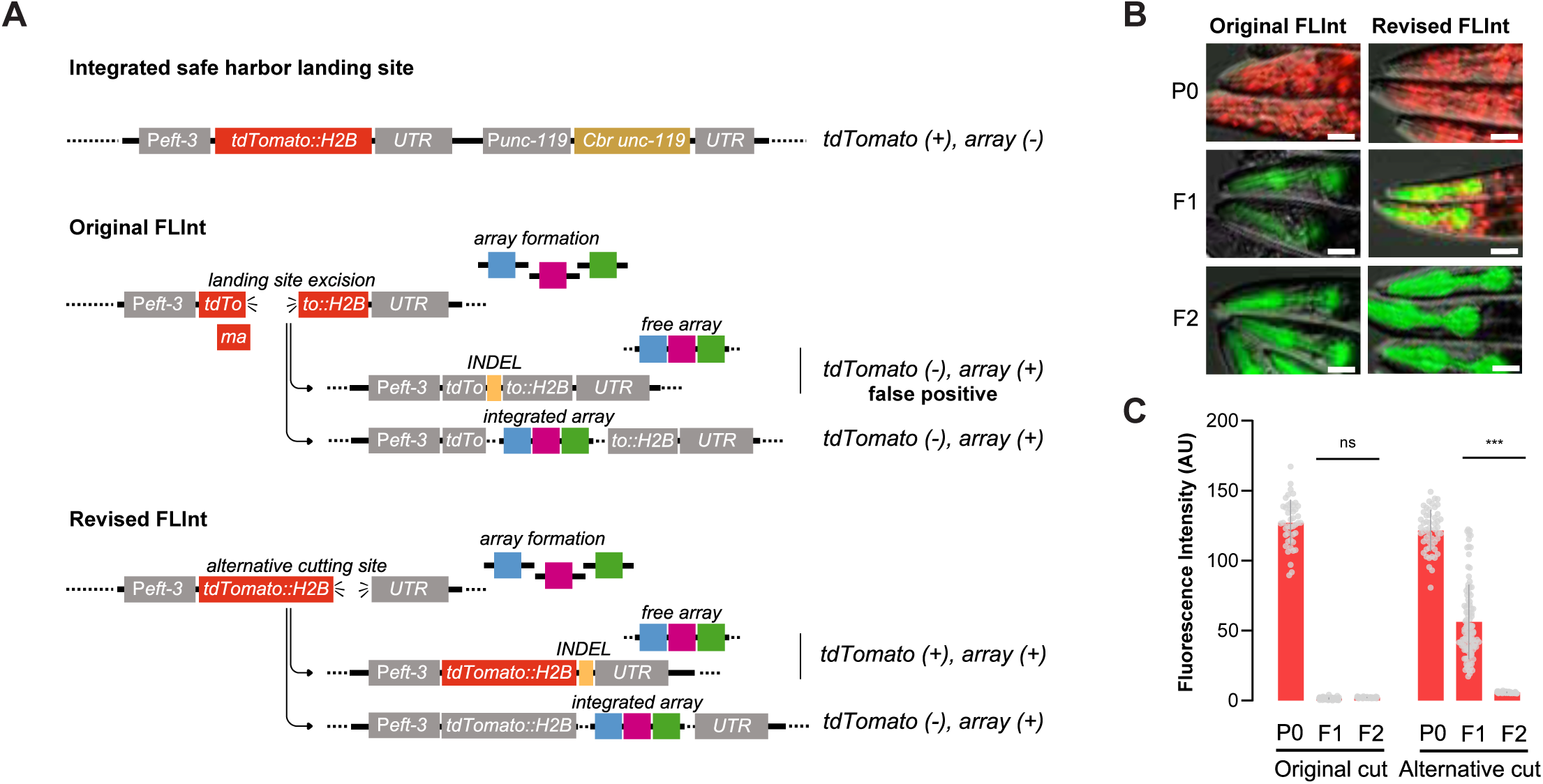
Revised FLInt enables phenotypic discrimination of true integrants. (A) Schematic of the original FLInt (FLInt 1.0) and revised FLInt (FLInt 2.0) strategies at the miniMos landing site (*eft-3p::tdTomato::H2B::unc-54 3′UTR; Cbr-unc-119(+)*). In FLInt 1.0, Cas9 cutting within the tdTomato coding region frequently generates INDEL alleles that abolish tdTomato expression, producing tdTomato-negative false positives. In FLInt 2.0, the cut site is shifted toward the 3′ region, allowing most INDEL alleles to retain tdTomato expression, while array integration disrupts tdTomato. (B) Representative fluorescence images of P0, F1, and F2 animals from the original FLInt and revised FLInt (20X objective lens) showing green pharyngeal co-injection marker and tdTomato nuclei. (C) Whole-animal tdTomato fluorescence intensity quantified from the images in (B) by measuring the mean red-channel signal using a segmented line region of interest in ImageJ. Sample sizes were as follows: Original cut P0 (n = 44), Original cut F1 (n = 91), Original cut F2 (n = 57), Alternative cut P0 (n = 51), Alternative cut F1 (n = 67), and Alternative cut F2 (n = 62). Data are shown as individual animals with mean ± SD. Statistical comparisons were performed using an unpaired *t*-test. ns, not significant; ***p < 0.001.

We first targeted the end of the H2B region, just before the 3’ UTR (3’ CDS, Fig 1A). Because CRISPR-induced double-strand breaks (DSBs) in *C. elegans* are frequently repaired by non-homologous end joining (NHEJ), microhomology-mediated end joining (Friedland et al. 2013), or synthesis-dependent strand annealing (Paix et al. 2017; Farboud et al. 2019), we expected that random indels would preserve a functional portion of tdTomato::H2B and maintain a functional 3′ UTR, restoring mRNA integrity and fluorescence despite loss of a CRISPR/Cas9 cutting site. To reduce edits in the coding region or tdTomato::H2B, we designed our crRNA on the antisense strand, which is predicted to result in indels predominantly 5’ to the CRISPR cut site, which is 3’ relative to the tdTomato::H2B reading frame (Farboud et al. 2019). Unless otherwise indicated, for all injections, we included a *myo-2*p::GFP; *rps-0*p::hygR co-marker plasmid (pMOD248) to mark array-positive animals for visual and antibiotic selection (Table 1). Three days post-injection, we screened F1 progeny showing pharyngeal GFP expression (indicative of array formation) and measured tdTomato fluorescence intensity (indicating potential presence of CRISPR-generated edits). Due to high efficiency of CRISPR-Cas9 relative to extrachromosomal array formation, we assumed that most array-positive (GFP+) worms bore chromosomal edits, which was supported by amplicon sequencing of pooled or single F1 and F2 GFP+ worms (Fig. 1B, Supplementary Fig. S1).

Thus, we considered GFP+ F1 animals likely to be a representative population of CRISPR-induced end-joining repair events at a given crRNA cut site. Compared with F1 progeny from the original FLInt 1.0 configuration, indel-bearing animals edited at the 3’ CDS exhibited higher tdTomato intensities, suggesting that these sites indeed promote selective fluorescence retention after CRISPR-mediated cutting (Fig. 1C, Fig. 2A-B). In contrast, F2 animals bearing chromosomally-integrated plasmids appeared to completely lack tdTomato expression in both approaches (Fig. 1C). These data suggest that by distinguishing true positive integrants from false positives via maintenance of fluorescence in the latter case, multi-copy integrants can likely be efficiently identified using this method.

**Figure 2.**
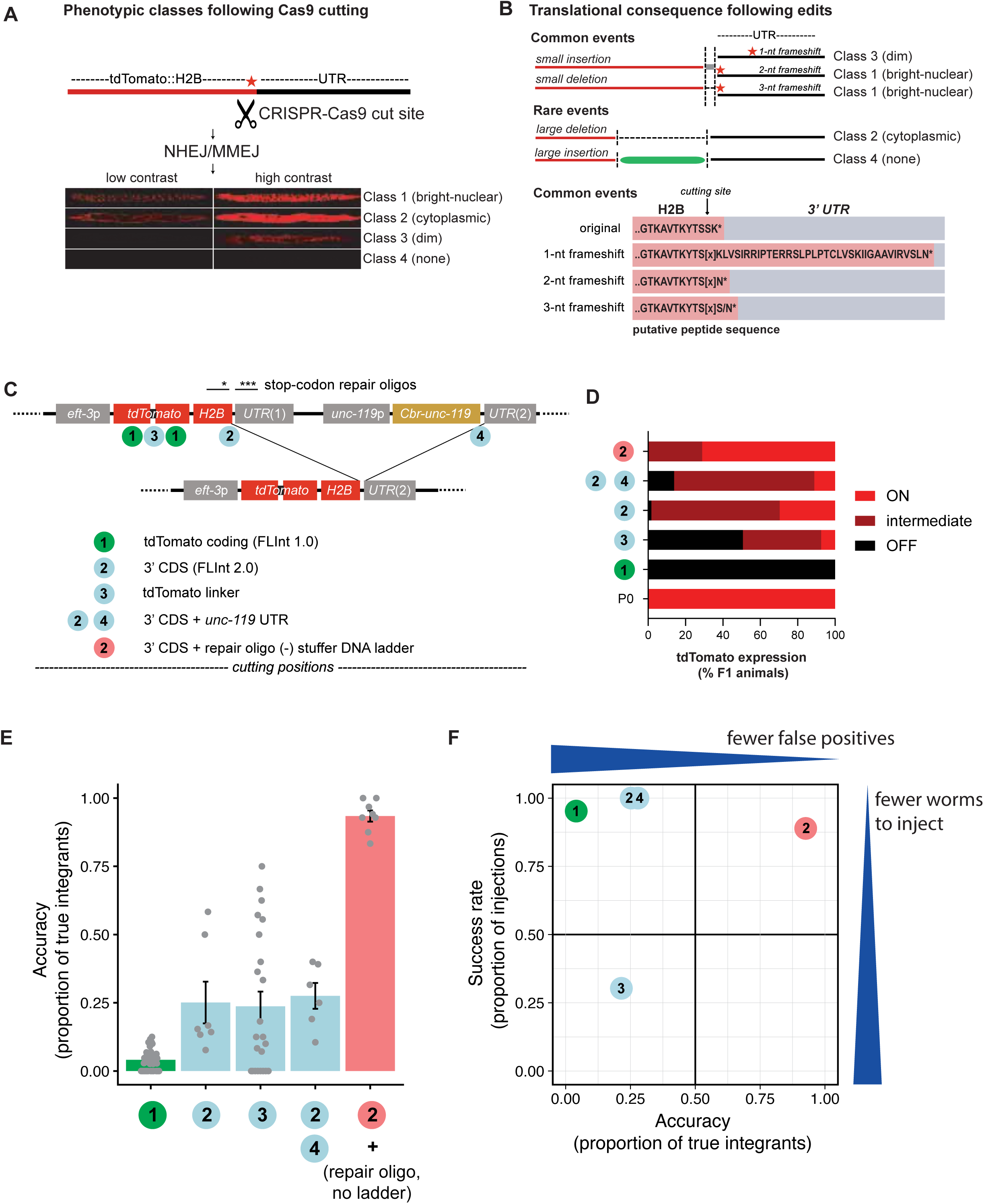
Cutting-site geometry determines tdTomato repair outcomes and integration accuracy. (A) Schematic of tdTomato::H2B::UTR showing the original Cas9 cut site and stop codon, followed by repair through non-homologous end joining or microhomology-mediated end joining (NHEJ/MMEJ). Representative tdTomato phenotypes are classified into four categories: class 1 (bright nuclear), class 2 (cytoplasmic), class 3 (dim), and class 4 (no signal), displayed at low and high contrast. (B) Predicted translational consequences of different repair outcomes. Small insertions or deletions generating 2-3 nt frameshifts preserve tdTomato expression via alternative stop codons (classes 1 and 3), whereas large insertions or deletions disrupt tdTomato localization or expression (classes 2 and 4). (C) Schematic of alternative CRISPR cutting strategies at the tdTomato::H2B::UTR–*unc-119*p::Cbr-*unc-119*::UTR locus. Cutting sites include tdTomato coding sequence (site 1, FLInt 1.0), H2B 3′ coding sequence (site 2, FLInt 2.0), tdTomato linker (site 3), dual cuts at H2B 3′ CDS and unc-119 UTR (sites 2+4), and site 2 with repair oligos without stuffer DNA. (D) Proportions of tdTomato expression phenotypes (ON, intermediate, OFF) among F1 progeny selected by the *myo-2*p::GFP co-injection marker for each cutting strategy. (E) Integration accuracy, defined as the proportion of selected candidates that were confirmed true integrants, across cutting strategies. Individual data points represent independent injections; bars indicate mean ± SEM. (F) Relationship between integration accuracy and success rate (proportion of successful injections), illustrating trade-offs between false-positive reduction and experimental efficiency.

### An intact tdTomato::H2B and 3′ UTR is essential for maintenance of fluorescence in non-integrated animals

While maintenance of fluorescence expression using a 3’ CDS crRNA improved the fraction of false-positive animals (those bearing low/no tdTomato expression but without an integrated transgene), there nonetheless appeared to be several different classes of fluorophore expression phenotypes generated, potentially confounding the identification of true positives. To better understand the molecular basis of INDEL-mediated maintenance of tdTomato expression, we examined the nature of repair among animals that did not carry integrated arrays. From injections targeting the 3’ CDS, we selected four classes of transgenic F2 animals based on their distinct tdTomato expression phenotypes: (1) bright nuclear tdTomato, (2) cytoplasmic tdTomato, (3) dim tdTomato, and (4) non-tdTomato (Fig. 2A). Genomic DNA from animals of each category was extracted, and the CRISPR-targeted region encompassing the H2B–UTR junction was amplified via PCR and sequenced (Fig. 2B). All sequenced animals carried edits at the intended cut site, confirming that CRISPR-mediated cutting is highly efficient among array-positive animals. Edited class 1 alleles (high tdTomato expression, n=8) restored either an in-frame junction between H2B and the *unc-54* 3’ UTR, or resulted in frameshift mutations followed almost immediately by an alternative stop codon in the 3’ UTR – consistent with functional re-coupling that stabilizes mRNA and maintains nuclear fluorescence (class 1 alleles, Fig. 2A-B). A second, rare class of alleles (class 2, n=7, Fig. 2A-B) bore larger deletions in the H2B coding sequence. Class 2 alleles exhibited bright, cytoplasmic tdTomato in edited worms. Both of these first two classes of alleles were expected and are easily distinguished from true-positive integrants. However, the third class of tdTomato expression alleles (dim class 3, n=9, Fig 2A-B) indicated the presence of frameshift mutations followed by long stretches of amino acids prior to a stop codon, potentially destabilizing the fluorescent protein. This suggests that maintenance of stop codons in all three reading frames proximal to the cut site might be important to avoid false positives in this configuration. We hypothesized that these alleles could be eliminated by modifying the editing process via inclusion of oligonucleotide repair templates intended to introduce stop codons in all three reading frames in addition to the existing stop codons in the 3’UTR. Consistent with this idea, inclusion of stop-codon-bearing oligonucleotide repair donors largely eliminated class III dimly fluorescent worms (Fig. 2C-D).

Because a larger number of F1 progeny generated by 3’CDS cutting retained tdTomato fluorescence compared to the previous method, we expected that only array-integrated lines would lose tdTomato expression in the subsequent generation, once homozygosity was achieved. We reasoned that array insertion disrupts the integrity of the tdTomato::H2B transgene by decoupling its coding sequence from the 3′ UTR, thereby disrupting translation and/or destabilizing the resulting protein specifically in true integrants. To directly test this, we again performed CRISPR targeting of the H2B 3’CDS site in the tdTomato background strain EG7944, co-injecting the *myo-2*p::GFP + *rps-0*p::hygR plasmid for visual and drug selection with and without oligo repair donors. Three days after injection, F1 progeny were treated with hygromycin to select edited animals. Six days post-injection, we screened F2 progeny for two key features: (1) complete loss of tdTomato fluorescence, and (2) presence of GFP expression. Worms meeting both criteria were singled to individual NGM/OP50 plates. After incubation at 25 °C for three days, F3 progenies from each line were examined for 100% transmission of the array marker to confirm stable genomic integration. From F2 candidates injected with a 3’CDS crRNA without oligo-repair donors, approximately 20% represented true integrants, as verified by complete marker segregation (Fig. 2E-F). Compared to the original FLInt method, this represented a roughly five-fold increase in candidate selection accuracy (Fig. 2E). There remained, however, a significant proportion (∼80%) of class-4 (tdTomato-negative) isolates that did not appear to harbor stable plasmid integration events, as measured by 100% transmission of the *myo-2*p::GFP (Fig. 2E). We were unable to obtain PCR amplicons of the safe-harbor locus from any of these isolates (n = 9), suggesting that these animals likely bore large, unintended insertions or deletions.

The majority of these false positive animals likely contained extrachromosomal arrays. Complex extrachromosomal arrays – those containing a complement of heterogeneous DNA sequences – are known to be more stably expressed than simple homogeneous sequences (Kelly et al. 1997), however *de novo* formation and inheritance of extrachromosomal arrays does not simply rely on DNA complexity *per se*, but rather an interaction between DNA AT-base composition and unknown other factors (Lin et al. 2021). We reasoned that elimination of carrier DNA might bias toward the inheritance of integrated arrays over extrachromosomal arrays. As expected, injections of low concentrations of *myo-2*p::*gfp* plasmid with and without carrier DNA both resulted in identification of GFP-positive, non-integrated F1 animals, but inheritance beyond F1 required the coinjection of carrier DNA (Table S1). Elimination of carrier DNA and inclusion of oligo HDR donors further increased the true-positive FLInt identification accuracy to nearly 100% (Fig. 2E-F), indicating that some false positives were likely due to either class 3 frameshift mutations at the chromosomal locus or due to unintended large indels (class 4) in animals also carrying extrachromosomal arrays (Fig. 2B). Because the accuracy of integrant selection at this stage is much higher than in the original FLInt method, this corresponds a dramatic reduction in the screening workload required to identify true integrants.

### FLInt 2.0 results in highly efficient multi-copy transgene integration with a simple workflow

Mechanisms of DNA repair vary depending on the nature of the DSB as well as the types of donor DNA available for repair (Paix et al. 2017; Farboud et al. 2019). We considered the possibility that larger deletions induced by multiple DSBs might alter the efficiency of integration of large, multi-copy arrays. Additionally, excision of a larger region of the safe harbor site offers the possibility to eliminate expression of the *C. briggsae unc-119+* rescue cassette, which may lead to pleiotropic effects in certain circumstances (Omi and Pujol 2021). We designed crRNAs targeting different positions within the same background strain that are predicted to restore fluorescence in the absence of frameshift mutations after editing by linking tdTomato::H2B to a downstream 3’UTR (Fig. 2C). To avoid generating uncoordinated worms, we outcrossed the existing *unc-119(ed3)* mutation prior to injections (Frøkjær-Jensen et al. 2014). Excision of this larger fragment resulted in a similar proportion of dimly fluorescent worms in the absence of integration, indicating that frameshift mutations are also likely responsible for this class of worms in this alternative configuration (Fig. 2D). Integration with coincident deletion of the *Cbr-unc119* coding sequence did not substantially increase selection accuracy relative to the single 3’ CDS crRNA indicating that a single DSB is sufficient for high efficiency integration in these conditions (Fig. 2E). In contrast, a crRNA targeting the linker domain between the two tdTomato dimers resulted in a larger proportion of dim, but still visible tdTomato-expressing offspring (Fig. 2C-D). Taken together, these results suggest that targeted insertion to the 3’ CDS of a fluorescent protein largely eliminates frameshift-induced false positive integrant identification via FLInt and that recoupling to a functional 3’ UTR enhances screening accuracy.

While selection accuracy provides a useful guide for choosing crRNA sites via FLInt, it does not necessarily correlate with overall success rate, as defined by the probability of identifying an integrant from a given injection batch. For example, despite generating a large number of dimly fluorescent F1 offspring using a tdTomato linker crRNA, the selection accuracy of true integrants using this configuration was similar to that obtained using the 3’ CDS crRNA (Fig. 2E). This is likely because the dimly fluorescent offspring are possible, yet difficult, to distinguish from an absence of fluorescence in these worms. However, identification of true integrants using a linker crRNA required the injection of many more P0 worms per batch, possibly because HDR repair in *cis* is highly efficient and may compete with, or occur prior to, transgene integration (Fig. 2F, Supplemental Fig. 1C). We compared the probability of integrant isolation for each of the crRNA cutting sites on a per-batch basis over many injections to estimate the efficiency of transgene insertion (Fig. 2F). This analysis shows that the 3’ CDS method using HDR donors is both the most accurate and most efficient, typically requiring fewer than 15 injected worms to reliably identify multiple integrants. This efficiency is comparable to the generation of extrachromosomal arrays in our hands.

### Copy number control via backbone excision

Previously described approaches for transgene integration indicated that transgene copy number is often correlated with expression level, which is apparent when a transgene contains a fluorescent coexpression marker (Yoshina et al. 2016; Noma and Jin 2018; Malaiwong et al. 2023). Preliminary examination of fluorescence levels in integrated transgenic lines generated using our method indicated that, as in the case of FLInt 1.0, these likely reflected high copy number transgenes. qPCR analysis of GFP copy number in 20 independent lines derived from our different integration methods indicated that all contained > 30-100+ copies, with no obvious effect of the cutting site on copy number (Supplemental Fig. 1D). While high-copy transgenes are preferable in many experimental contexts, it is often desirable to tailor copy number and expression level, which can be a laborious process using standard methods. We reasoned that we could selectively reduce transgene copy number simply via CRISPR-mediated excision.

We designed crRNAs which are predicted to cut within the *beta-lactamase* (*bla*/*ampR)* gene contained in many common cloning vectors (Table 1). Cas9 targeting of this site via crRNA injection in a high-copy number transgene line resulted in highly efficient generation of offspring with various levels of reduced expression of *myo-2*p::GFP (Fig. 3A). Some of these lines exhibited expression levels approximating that of a single-copy insertion (Fig. 3A). To determine whether *myo-2*::GFP serves as a reliable proxy for the coexpression level of additional transgenes, we first generated multi-copy integrated lines via FLInt 2.0 that coexpressed mCherry in the CAN cell, via *ceh-23p* (Wenick and Hobert 2004, Table 1). Copy number reduction via *ampR* crRNA injection in these lines lead to a consistent, relative reduction in expression of both GFP and mCherry, indicating that *myo-*2::GFP reliably predicted the expression of array-encoded transgenes under these conditions (Fig. 3B).

**Figure 3.**
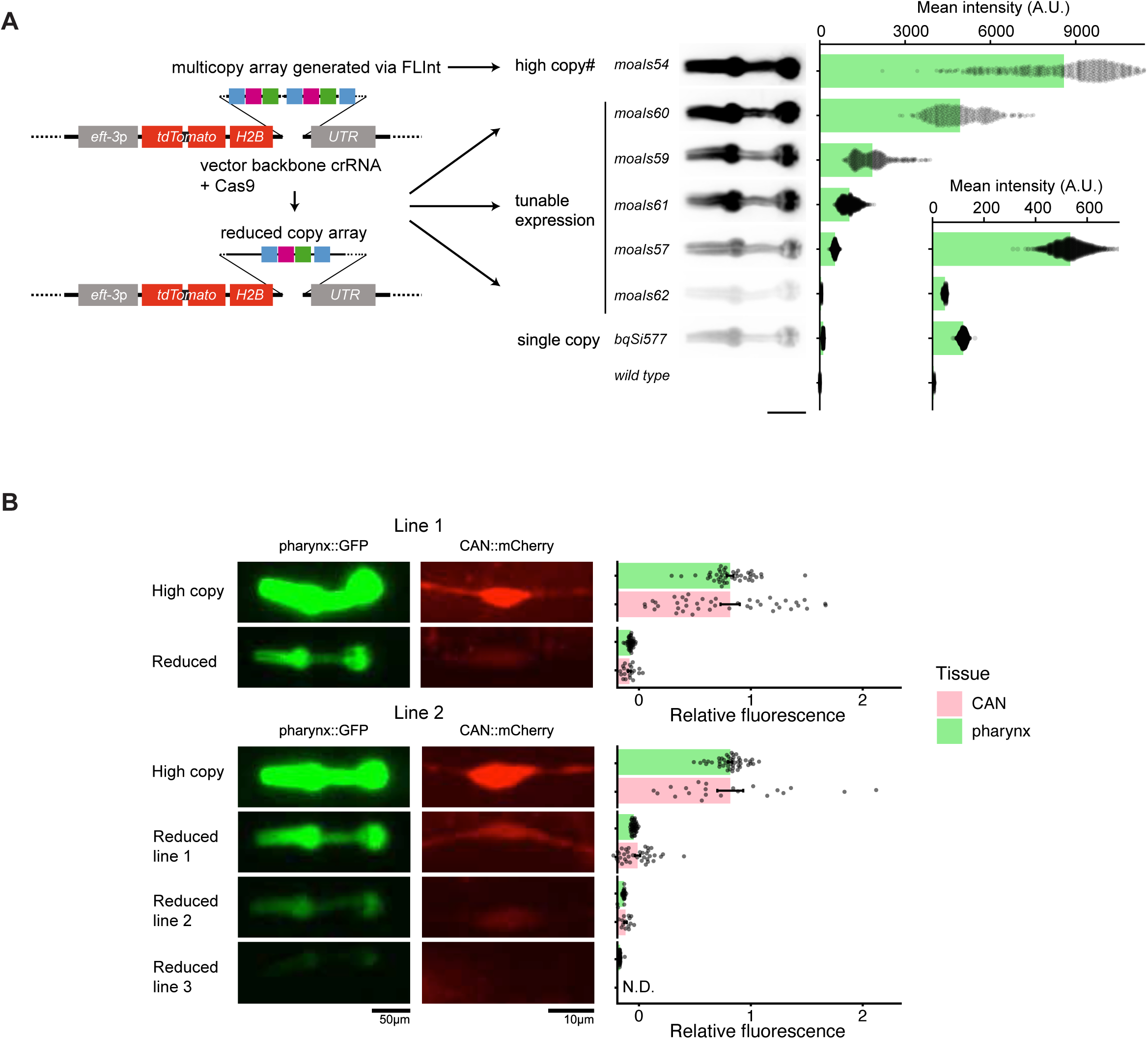
FLInt enables tunable transgene expression by reducing extrachromosomal array copy number. (A) Left, schematic illustrating conversion of a high-copy extrachromosomal array generated by FLInt into a reduced-copy array through CRISPR–Cas9 cleavage of the vector backbone using a backbone-specific crRNA. Targeting the backbone decreases array copy number without disrupting the *eft-3p::tdTomato::H2B* expression cassette, resulting in tunable expression levels. Representative fluorescence images of independent lines with varying copy numbers (*moals54* (background strain), *moals60*, *moals59*, *moals61*, *moals57*, *moals62*, and *bqSi577* (single-copy via MosSCI) are shown alongside wild-type controls. Right, quantification of tdTomato fluorescence intensity (A.U.) demonstrates a graded reduction in expression corresponding to decreasing array copy number. Each dot represents the average fluorescence of a region of interest in an individual animal. Scale bar = 100 µm. (B) Correlation of expression of pharyngeal GFP and CAN mCherry in high copy and reduced copy transgenic lines after targeting vector backbone sequence with *ampR* crRNA. Upper and lower panel represent copy number reduction technique using two independently derived multicopy array lines, respectively. Each dot represents the average fluorescence of a region of interest in an individual animal divided by the population average expression for the respective tissue type in the multi-copy parental strain. Bars indicate mean; error bars are s.e.m. Scale bars are indicated. N.D., not detectable.

However, some plasmids have been reported to exhibit non-random distribution in integrated arrays (Rich et al. 2025), so this approach should be verified for each plasmid coinjection combination, or via inclusion of transgenes in the same vector. Previous experiments using FLInt 1.0 indicated that copy number correlated with the concentration of injected plasmids, but experimentally this required separate injections, increasing workload (Malaiwong et al. 2023). These results indicate that high-copy-number transgenes generated via FLInt can be easily tuned to desirable expression levels via excision using Cas9 and vector backbone targeting.

### FLInt 2.0 requires similar resources as generation of extrachromosomal arrays

A motivation for this work was to develop a method for efficient transgene integration requiring few or no specialized reagents that could replace extrachromosomal array generation. The cost of reagents (Cas9, crRNAs, oligos, plasmid, hygromycin) using this method are not prohibitive for most labs, when estimated per integrated line. However, an additional barrier to uptake of integration methods is the amount of effort, time and resources required to identify the desired strains. We therefore estimated the resources required to generate integrated strains in two different ways: hours of effort per integrant construct/line isolated and number of plates used. Based on reported efficiencies and in our experience, FLInt 1.0 and other comparable methods, including extrachromosomal array-based methods, all typically require the injection of ∼15-20 P0 animals per transgene and vary by experience, so we have excluded the injection time in our estimates (El Mouridi et al. 2022; Yoshina and Mitani 2022; Malaiwong et al. 2023; Yanagi and Lehrbach 2024). Because the selection process for FLInt 2.0 occurs in the F2 generation and animals are pooled after an injection session, this results in a significant reduction in time prior to screening, with only a few minutes required for addition of hygromycin 3 days after injection (Fig. 4A). While hygromycin selection can be omitted if desired, this significantly reduces the screening time required to identify candidates in the subsequent F2 stage, by eliminating offspring from poorly injected animals or non-transgenic offspring. Identification and isolation of 15-20 co-expression marker-positive and tdTomato-negative F2 worms typically takes 30 minutes. Screening of F3 animals to distinguish between extrachromosomal array lines, heterozygous and homozygous integrant lines requires an additional 30 minutes. Because this approach usually results in the isolation of multiple integrated lines, this approach typically yields several integrated transgene lines with about 2 hours of effort.

**Figure 4.**
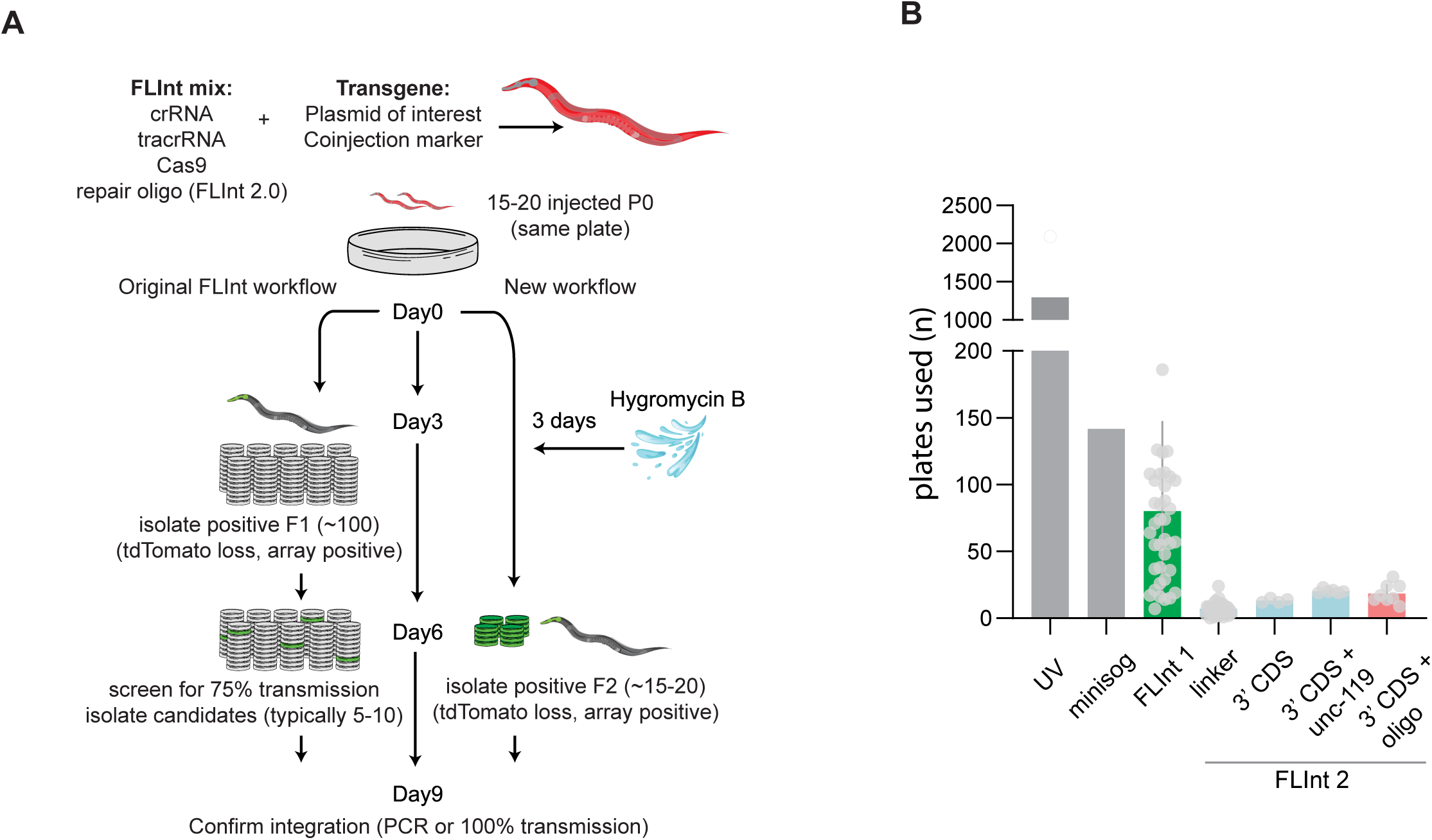
Comparison of FLInt workflows and plate usage across integration methods. (A) Schematic outlining the original and updated FLInt workflows, showing the composition of the FLInt injection mix, the number of injected P0 animals, and the stages at which F1 and F2 candidates are isolated based on tdTomato loss and array positivity, followed by confirmation of integration by PCR or transmission analysis. (B) Quantification of the number of plates used for isolating integrants across different integration strategies, including UV (Mariol et al. 2013), miniSOG (Noma and Jin 2018), FLInt 1 (Malaiwong et al. 2023), FLInt linker, and FLInt 2 configurations (3′ CDS, 3′ CDS + *unc-119*, 3′ CDS + repair oligo). Each dot represents an independent experiment; bars indicate mean plate usage.

Screening for integrant strains using conventional methods typically requires the isolation of many candidate lines, followed by segregation analysis over multiple generations, resulting in the use of substantial quantities of materials (Fig. 4A-B). In contrast, this method requires only about 15-20 plates in total prior to integrant identification (Fig. 4B). Together, these results suggest that the combination of effort required and materials used in this method approximates that of stably-inherited extrachromosomal array line generation.

## Discussion

Here we describe a simple and accessible method for the generation of multi-copy transgenes in *C. elegans.* Our method eliminates existing barriers to the adoption of multi-copy integrated transgenes over extrachromosomal arrays; this approach is affordable for most labs based on cost and effort and requires no specialized cloning or reagents other than Cas9 ribonucleoprotein complex.

### Current approaches to generation of integrated transgenes

Recent advances in genome engineering have made the introduction of precise edits and insertions in *C. elegans* using CRISPR-Cas9 highly efficient (Dickinson and Goldstein 2016; Nance and Frøkjær-Jensen 2019). Similarly, the development of safe harbor landing sites in the genome via Mos1 (Frøkjaer-Jensen et al. 2008; Frøkjær-Jensen et al. 2012; Frøkjær-Jensen et al. 2014; El Mouridi et al. 2022) has enabled precise and efficient single copy transgene insertion via FLP recombinase (Nonet 2020; Nonet 2023), CRISPR-Cas9 (Norris et al. 2015; Philip et al. 2019; Silva-García et al. 2019; Stevenson et al. 2020; El Mouridi et al. 2022) or PhiC31 integrase (Yang et al. 2022). In contrast, there are relatively few developed methods to efficiently integrate large multi-copy or multi-plasmid arrays in defined genomic loci, which may contribute to the continued use of extrachromosomal arrays in many labs when high levels of expression or the expression of multiple constructs is needed. *C. elegans’* natural capacity to readily generate complex, recombinant megabase-sized extrachromosomal arrays of foreign DNA with holocentric properties remains a unique strength among experimental organisms (Stinchcomb et al. 1985; Mello et al. 1991; Lin et al. 2021; Stevenson et al. 2023), yet inheritance of these arrays is meiotically and mitotically unstable and they are frequently silenced in the germline, limiting their robust experimental application.

Recently, El Mouridi and colleagues developed a powerful method to integrate existing complex extrachromosomal arrays using CRISPR-Cas9-dependent excision and non-homologous repair at safe harbor landing sites (El Mouridi et al. 2022). These efforts have been complemented by methods to integrate arrays in a single shot injection, via integration at CRISPR-Cas9-generated DSBs in the *dpy-3* or *ben-1* genes (Yoshina and Mitani 2022) or in safe harbor landing sites (Malaiwong et al. 2023; Yanagi and Lehrbach 2024) or via PhiC31 integrase mediated integration at defined loci (Rich et al. 2025). While all of these methods have strengths, they either require multiple injection steps or result in a non-optimal proportion of false positives, increasing screening time and effort. Here, we sought to develop methods that are comparably as tractable as extrachromosomal array construction but that obviate the instability of these arrays.

### Simplicity and generalizability of this system

The method we describe here requires no cloning, is theoretically compatible with any plasmid or PCR product and can be used to generate multi-copy arrays from a single injection of ∼15 animals with very little screening effort and few false positives. We have shown that arrays generated using this method typically contain high copy numbers, as was the case with the previous FLInt method (Malaiwong et al. 2023). By coinjection with a readily visible or selectable transgene marker (e.g. *myo-2*p::*gfp*, *hygR*), screening time using this approach requires minutes, not hours, using a fluorescent stereomicroscope. We also show that copy number can be readily tailored to desired expression level using a simple additional CRISPR-Cas9 targeting of common sequences included in cloning vectors, likely eliminating the need for laborious injections of variable construct concentrations. A detailed protocol is actively maintained at the following URL: dx.doi.org/10.17504/protocols.io.eq2lyqm1pvx9/v1.

### Future extensions

In our experiments, we have used FLInt 2.0 to generate relatively simple multi-copy arrays (1-4 plasmids, with or without “stuffer DNA” ladder). While additional work is required to establish the upper limit for transgene complexity using this method, we anticipate that this should in theory approach that of highly complex extrachromosomal arrays analyzed in other work (Lin et al. 2021; El Mouridi et al. 2022; Stevenson et al. 2023). This suggests that integrated transgene library assembly should be readily tractable using this approach or combined with other existing approaches.

## Data availability

Strains and plasmids are available upon request. The authors affirm that all data necessary for confirming the conclusions of the article are present within the article, figures, and tables.

## Acknowledgements

We thank members of the O’Donnell lab for thoughtful discussions of the manuscript. Some strains were provided by the CGC, which is funded by NIH Office of Research Infrastructure Programs (P40 OD010440).

## Study funding

This work was funded via the NIH (DP2 GM154014 to M.P.O).

## Conflict of interest

The authors declare no conflict of interest.

